# Three-dimensional object-based quantification of nicotinic acetylcholine receptor subunit colocalization using Imaris

**DOI:** 10.64898/2026.10.11.758376

**Authors:** Ankura Sitaula, Aref Zarin

## Abstract

Neural circuit function depends not only on synaptic connectivity but also on the molecular composition and spatial organization of neurotransmitter receptors. Determining whether receptor subunits occupy shared or distinct domains requires spatial measurements capable of resolving sparse receptor puncta across neural tissue and, when needed, restricting analysis to defined cellular compartments. Conventional intensity-based colocalization metrics can be difficult to interpret for sparse, punctate signals and do not inherently provide object-level classification or cell-specific analysis. Here, we describe a three-dimensional object-based colocalization workflow for endogenously tagged nicotinic acetylcholine receptor (nAChR) subunits imaged by Airyscan super-resolution confocal microscopy and analyzed in Imaris. Individual receptor puncta are segmented as 3D surface objects and classified as colocalized or non-colocalized using a control-defined spatial overlap criterion. The workflow supports analysis within a defined region of the ventral nerve cord neuropil or within genetically labeled motor-neuron dendritic compartments. For dendrite-restricted analysis, receptor puncta are first selected by their overlap with the dendritic surface before receptor–receptor overlap is quantified. The overlap criterion is empirically calibrated using positive and negative biological controls processed under the same imaging and analysis conditions, providing a reproducible alternative to arbitrary threshold selection. This approach enables receptor organization to be quantified within the neuropil and genetically labeled motor-neuron dendritic compartments and can be adapted to other epitope-tagged receptors, ion channels, and synaptic proteins.

**Key features:**

- Enables 3D object-based colocalization analysis of sparse, punctate receptor signals that may be difficult to evaluate using conventional intensity-based approaches.
- Supports colocalization analysis at two spatial scales: within defined neural tissue regions and within genetically labeled neuronal dendritic compartments.
- Particularly suited for endogenously tagged receptors and other punctate membrane proteins imaged by high-resolution confocal or Airyscan microscopy.
- Uses experimentally validated positive and negative controls to establish overlap criteria for classifying individual 3D puncta as colocalized or non-colocalized.

**This protocol is used in:** PNAS Nexus (2026), DOI: https://doi.org/10.1093/pnasnexus/pgag173

**Graphical overview:** **Graphical overview. 3D object-based workflow for analyzing nAChR subunit colocalization in the *Drosophila* larval ventral nerve cord**. (A) Dissect and immunostain third-instar larvae, then acquire a three-dimensional (3D) Airyscan image stack of the selected ventral nerve cord (VNC) region. (B) Segment the fluorescence signals of receptor subunits X and Y as separate 3D surfaces in Imaris. (C) Define the overlap criterion using a positive control (dual -tagged α6::HA and α6::OLLAS; >98% overlap) and a negative control (Rdl::HA and α5::OLLAS; <2% overlap). Classify objects as overlapping when the *Overlapped Volume to Surfaces* value is ≥0.05 µm^3^ . (D) Assess receptor overlap using one of two analysis regions: the dorsal VNC neuropil or motor-neuron (MN) dendrites. For the MN dendrite-restricted analysis, first retain the surfaces of each receptor subunit that overlap the labeled dendrites, then assess overlap between the two retained receptor populations. (E) Count overlapping (CoLo) and non-overlapping (N.CoLo) receptor objects. The bar graph illustrates the two output categories; its values do not represent experimental data.

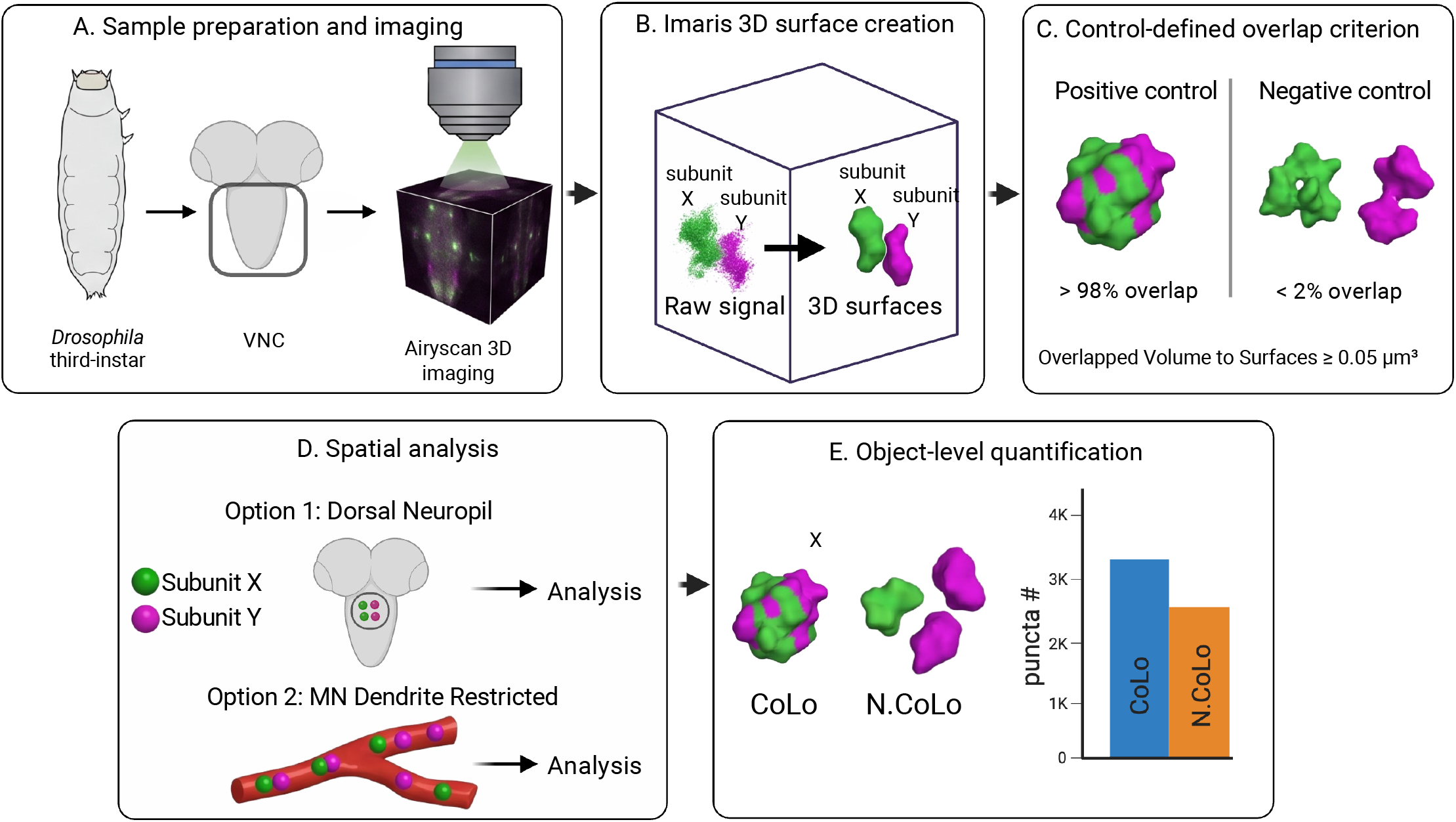

## Background

The function of a neural circuit depends not only on which neurons are connected but also on the molecular composition of their synapses. In particular, the identity and subcellular localization of neurotransmitter receptors influence how individual neurons respond to synaptic input. Advances in electron microscopy have enabled neural circuits to be reconstructed at synaptic resolution, but anatomical connectivity alone does not reveal the molecular composition of postsynaptic sites. Recent studies in *Drosophila* have therefore begun combining connectomic information with endogenous receptor labeling to map neurotransmitter receptors within identified neurons and dendritic compartments [1-3]. Such studies have revealed that receptor subtypes can exhibit highly specific subcellular distributions, providing an additional molecular layer of organization within anatomically defined circuits.

nAChRs provide an excellent system for investigating this organization. nAChRs mediate fast excitatory cholinergic transmission in the *Drosophila* central nervous system and are assembled from a diverse family of receptor subunits [4,5]. A defining feature of nAChRs is their molecular complexity, which gives rise to substantial structural and functional diversity. The *Drosophila* genome encodes 10 nAChR subunits—seven α and three β—which can assemble into different pentameric receptor complexes with distinct functional properties [4,6,7]. In our recent study, we identified at least eight nAChR subunits expressed in larval motor neurons (MNs) and found that different subunit pairs exhibit distinct spatial relationships [3]. These findings raise an important methodological question: how can the spatial association between punctate receptor populations be quantified reproducibly, particularly when the analysis must be restricted to a defined neural region or an identified neuronal compartment?

Fluorescence colocalization can be evaluated using several analytical strategies. Common intensity -based methods assess relationships between fluorescence intensities across pixels or voxels, whereas object-based methods first segment fluorescent structures into discrete objects and subsequently evaluate their spatial relationships [8,9]. Different approaches have distinct advantages and limitations, and the appropriate method depends strongly on the biological question and the spatial characteristics of the fluorescent signal [8,9]. Although intensity-based analysis can be valuable, threshold selection is often subjective, and measured intensities may be influenced by autofluorescence, spectral bleed-through, background noise, and variation in image acquisition. Moreover, these methods quantify signal co-occurrence or covariance at the pixel or voxel level rather than within biologically defined structures. Statistical procedures have also been developed to establish objective thresholds and evaluate whether measured colocalization exceeds that expected from random spatial association [10]. For sparse receptor labeling that can be resolved as individual puncta, object-based analysis is particularly useful because each punctum can be analyzed as an individual spatial object and its relationship with another receptor population can be evaluated directly.

Here, we describe a three-dimensional object-based workflow developed to quantify the spatial organization of endogenously tagged nAChR subunits in the *Drosophila* larval VNC. Airyscan super-resolution image stacks are analyzed in Imaris, where receptor puncta are segmented as 3D surface objects. The protocol supports two complementary levels of analysis. First, receptor–receptor colocalization can be quantified within a defined region of the VNC neuropil. Second, a genetically labeled MN dendritic compartment can be segmented as an additional 3D surface, allowing receptor puncta to be selected based on their association with the MN before receptor–receptor colocalization is quantified. These approaches were used in our original study to distinguish receptor organization across the dorsal neuropil from receptor organization specifically within MN dendrites [3].

An important feature of the workflow is the use of biological positive and negative controls to empirically establish the overlap criterion used to classify receptor puncta. A positive control consisting of two different epitope tags associated with the same receptor subunit establishes the level of overlap expected from two labels reporting the same receptor population, whereas a receptor pair expected to occupy largely distinct spatial domains provides a negative reference. Applying the same image acquisition, surface segmentation, and overlap analysis to the controls and experimental samples provides a consistent framework for classifying individual receptor puncta as colocalized or non-colocalized.

The use of endogenous receptor tagging is also advantageous for studying subcellular receptor organization because receptor localization can be examined without relying on conventional transgenic receptor overexpression. Endogenous neurotransmitter-receptor tagging strategies have previously been developed in *Drosophila* [1], and more recent conditional epitope-tagging approaches enable endogenous receptor populations to be visualized within genetically defined cell types [2]. The workflow described here can therefore be used to investigate receptor organization both across neural tissue and within genetically defined neuronal compartments.

Colocalization in this protocol represents spatial overlap at the resolution of the imaging and segmentation workflow and should not be interpreted as evidence that two receptor subunits physically interact or assemble within the same receptor complex. Instead, the method measures the extent to which two populations of receptor puncta occupy overlapping spatial domains. Although developed for nAChR subunits in *Drosophila* MNs, the general workflow can be adapted to other fluorescently or epitope-tagged receptors, ion channels, synaptic proteins, or other punctate cellular structures for which appropriate imaging and biological controls are available.

## Materials and reagents

### Biological materials

1. *Drosophila* stocks
  a. MN driver: D42-Gal4; UAS-mCherry [3]
  b. α6-GFP11-3xHA [11]
  c. nAChRα6-KDRT-smGdP-10xOLLAS [2]
  d. nAChRα5-KDRT-smGdP-10xOLLAS [2]
  e. nAChRα5::EGFP [12]
  f. nAChRα3-KDRT-1xALFA [2]
  g. α1-GFP11-3xHA [11]
  h. nAChRα2::EGFP [12]
  i. nAChRα4::EGFP [12]
  j. nAChRα7-KDRT-1xALFA [2]

### Reagents

1. Normal donkey serum (Jackson ImmunoResearch Laboratories, catalog number: 017-000-121)
2. Normal goat serum (Jackson ImmunoResearch Laboratories, catalog number: 005-000-121)
3. Fluoromount-G (SouthernBiotech, catalog number: 0100-01)
4. Paraformaldehyde, 16% solution, EM grade (Electron Microscopy Sciences, catalog number: 15710)
5. Phosphate-buffered saline (PBS) (Fisher Scientific, catalog number: BP2944-100)
6. Triton X-100 (Fisher Scientific, catalog number: BP151-500)
7. Sodium chloride (Fisher Scientific, catalog number: BP358-1)
8. Potassium chloride (Fisher Scientific, catalog number: BP366-500)
9. Calcium chloride dihydrate (Fisher Scientific, catalog number: BP510-500)
10. Magnesium chloride hexahydrate (Fisher Scientific, catalog number: BP214-500)
11. Sodium bicarbonate (Fisher Scientific, catalog number: S233-500)
12. HEPES (Fisher Scientific, catalog number: BP310-100)
13. Sucrose (Thermo Scientific, catalog number: 177140010)
14. Trehalose Anhydrous (TCI, catalog number: 99-20-7)
15. Mouse anti-HA (1:1000, BioLegend, catalog number: 901516)
16. Rat anti-HA (1:200, SigmaAldrich, catalog number: 11867423001)
17. Rat anti-OLLAS (1:100, Novus Biologicals, catalog number: NBP106713H)
18. Mouse anti-ALFA (1:100, Thermo Fisher Scientific, catalog number: MA5-54241)
19. Chicken anti-GFP (1:500, Thermo Fisher Scientific, catalog number: A10262)
20. Mouse anti-GFP (1:100, Developmental Studies Hybridoma Bank, catalog number: AB_2617417)
21. Rabbit anti-GFP (1:500, Thermo Fisher Scientific, catalog number: A-11122)
22. Mouse anti-mCherry (1:200, Takara Bio, catalog number: 632543)
23. Rabbit anti-mCherry (1:200, Abcam, catalog number: EPR20579)
24. Atto 488 Alpaca anti-ALFA (1:500, Nanotag, catalog number: N1502)
25. Rabbit anti-V5 (1:500, Cell Signaling Technology, catalog number: 13202)
26. Alexa Fluor 488 Goat anti-Mouse (1:200, Thermo Fisher Scientific, catalog number: A-11029)
27. Alexa Fluor 488 Goat anti-Rabbit (1:200, Thermo Fisher Scientific, catalog number: A-11008)
28. Alexa Fluor 488 Donkey anti-Chicken (1:200, Thermo Fisher Scientific, catalog number: A78948)
29. Alexa Fluor 555 Goat anti-Mouse (1:200, Thermo Fisher Scientific, catalog number: A-21422)
30. Alexa Fluor 555 Goat anti-Rat (1:200, Thermo Fisher Scientific, catalog number: A-21434)
31. Alexa Fluor 555 Goat anti-Rabbit (1:200, Thermo Fisher Scientific, catalog number: A-21429)
32. Alexa Fluor 594 Goat anti-Rat (1:200, Thermo Fisher Scientific, catalog number: A-11007)
33. Alexa Fluor 594 Goat anti-Rabbit (1:200, Thermo Fisher Scientific, catalog number: A-11012)
34. Alexa Fluor 647 Goat anti-Rat (1:200, Thermo Fisher Scientific, catalog number: A-21247)
35. Alexa Fluor 647 Donkey anti-Mouse (1:200, Thermo Fisher Scientific, catalog number: A-31571)
36. Alexa Fluor 647 Donkey anti-Rabbit (1:200, Thermo Fisher Scientific, catalog number: A-31573)

### Solutions

1. PBST: 0.3% (v/v) Triton X-100 in 1× PBS (see Recipes)
2. HL3.1 hemolymph-like solution (see Recipes).
3. 4% PFA in 1× PBST (see Recipes)
4. Blocking solution: 1% normal donkey serum and 1% normal goat serum in PBST (see Recipes).

### Recipes

#### 1. PBST

| Reagent | Final concentration | Quantity for 1 L |
| --- | --- | --- |
| PBS tablets (Fisher BioReagents, BP2944-100) | 1× PBS | 5 tablets |
| Deionized water | — | 1 L |
| Triton X-100 | 0.3% (v/v) | 3 mL |

*Note: Dissolve five PBS tablets in 1 L deionized water. Add 3 mL Triton X-100 and mix thoroughly. Store at 4 °C.*

#### 2. HL3.1

| Reagent | Final concentration | Quantity for 1 L |
| --- | --- | --- |
| Sodium chloride (NaCl) | 70 mM | 4.09 g |
| Potassium chloride (KCl) | 5 mM | 0.37 g |
| Calcium chloride dihydrate (CaCl<sub>2</sub>·2H<sub>2</sub>O) | 1.5 mM | 0.2205 g |
| Magnesium chloride hexahydrate (MgCl<sub>2</sub>·6H<sub>2</sub>O) | 4 mM | 0.814 g |
| Sodium bicarbonate (NaHCO<sub>3</sub>) | 10 mM | 0.84 g |
| Trehalose | 5 mM | 1.89 g |
| Sucrose | 115 mM | 39.36 g |
| HEPES | 5 mM | 1.19 g |
| Deionized water | — | Bring to 1 L |

*Note: Dissolve the components in deionized water and bring the final volume to 1 L with deionized water. Store at 4 °C.*

#### 3. 4% PFA in 1× PBST

| Reagent | Final concentration | Quantity for 100 mL |
| --- | --- | --- |
| 16% paraformaldehyde solution, EM grade (Electron Microscopy Sciences, 15710) | 4% (w/v) | 25 mL |
| 1× PBST | — | 75 mL |

*Note: Combine 25 mL of 16% PFA solution with 75 mL PBST and mix gently. Prepare fresh on the day of use. Handle PFA using appropriate chemical-safety precautions.*

#### 4. Blocking solution

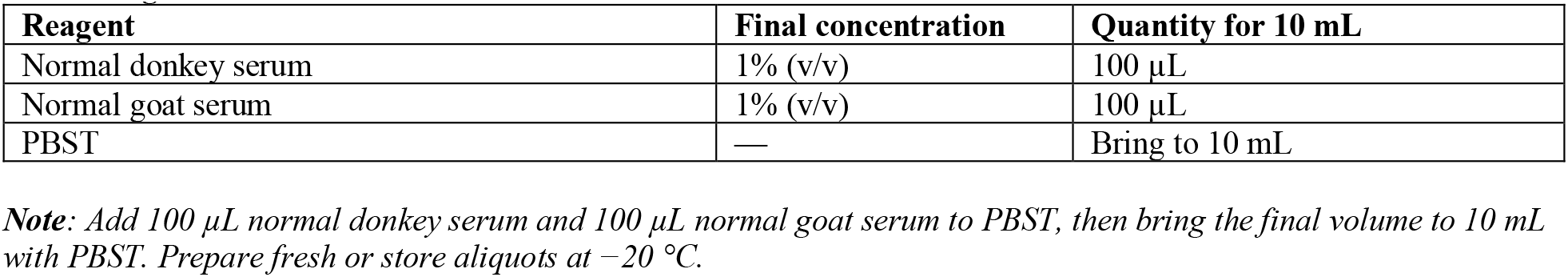

| Reagent | Final concentration | Quantity for 10 mL |
| --- | --- | --- |
| Normal donkey serum | 1% (v/v) | 100 µL |
| Normal goat serum | 1% (v/v) | 100 µL |
| PBST | — | Bring to 10 mL |

*Note: Add 100 µL normal donkey serum and 100 µL normal goat serum to PBST, then bring the final volume to 10 mL with PBST. Prepare fresh or store aliquots at −20 °C.*

### Laboratory supplies

1. 24-well plate (MilliporeSigma, catalog number: PSMT010R)
2. Forceps (MilliporeSigma, catalog number: F6521)
3. Petri dish 10 cm Ø (Sigma-Aldrich, catalog number: CLS430167)
4. Microscope slides (Fisher Scientific, catalog number: 12-550-400)
5. Coverslips (Fisher Scientific, catalog number: 12-547)

### Equipment

1. Zeiss LSM 900 confocal microscope
2. 63× oil-immersion objective (NA 1.4; mounted on the Zeiss LSM 900).
3. Stereomicroscope (ZEISS, model: Stemi 508)

### Software and datasets

1. ZEN Blue (ZEISS, version 3.7) was used for Airyscan image acquisition and three-dimensional Airyscan reconstruction. ZEN Blue requires an institutional or instrument-associated license. Available at: https://www.zeiss.com/microscopy/en/products/software/zeiss-zen.html (accessed September 21, 2026).
2. Fiji/ImageJ (version 1.54g) was used for channel visualization. Fiji is freely available open-source software under the GNU General Public License. Available at: https://imagej.net/imagej-wiki-static/Fiji (accessed September 21, 2026).
3. Imaris (Bitplane/Oxford Instruments, version 10.1) was used for three-dimensional surface segmentation and object–object colocalization analysis. Imaris requires an institutional or commercial license. Available at: https://imaris.oxinst.com (accessed September 21, 2026).
4. Airyscan image stacks and analysis code associated with this protocol are publicly available from Dryad: Sitaula, A.; Kaur, K. P.; Mogharrabi, A.; Olsen, L.; and Zarin, A. (2026). Data and code from: Motor neurons integrate cholinergic inputs through spatial organization of diverse nicotinic receptors. Dryad. https://doi.org/10.5061/dryad.fxpnvx17f (accessed September 21, 2026). The repository includes the Figure 5 receptor-colocalization image stacks and the Jupyter notebook used for colocalization analysis.

### Procedure

#### A. Sample preparation and immunostaining

1. Select third-instar larvae and dissect the VNC in HL3.1 hemolymph-like solution.
2. Fix samples in 4% paraformaldehyde prepared in 1× PBST for 10 min at room temperature in a 24-well plate.
3. Block samples overnight at 4 °C in PBST containing 1% normal donkey serum and 1% normal goat serum.
4. Incubate samples with the validated primary antibodies for the two endogenous nAChR tags and, when performing MN dendrite-restricted analysis, the dendritic marker. Incubate overnight at 4 °C.
5. Wash three times in PBST for 15 min per wash.
6. Incubate samples with the corresponding Alexa Fluor secondary antibodies at 1:200 overnight at 4 °C.
7. Wash three times in PBST for 15 min per wash, mount in Fluoromount-G, and store at −20 °C until imaging.

***Note:*** *Use antibody/tag combinations appropriate to the receptor pair. Primary antibodies and working dilutions are listed in Materials and reagents*.

#### B. Airyscan acquisition and 3D reconstruction

1. Place the mounted VNC slide on a Zeiss LSM 900 confocal microscope fitted with a 63× oil immersion objective (NA 1.4). Use Airyscan super-resolution mode in ZEN Blue to acquire image stacks from third-instar larvae.
2. Set the acquisition parameters for the fluorophores and objective available on the microscope. Optimize laser power, detector settings, zoom, and the z-stack to obtain receptor puncta that are clearly resolved without obvious saturation or loss of weak signal. These settings are experiment-dependent and therefore should be optimized locally rather than copied as fixed values.
3. Once the acquisition settings are selected for an experiment, keep the imaging parameters constant across all images being compared. Use the same objective, zoom, laser power, detector settings, scan settings, and channel configuration for samples intended for the same quantitative comparison.
4. Acquire a z-stack encompassing the VNC region required for the selected workflow. Save individual fluorescence channels separately and retain the original raw image stack.
5. Process each raw Airyscan z-stack in ZEN Blue using 3D Airyscan processing with the automatic settings. Save the reconstructed stack with its channel identities preserved; use this reconstructed stack for subsequent analysis in Imaris Bitplane 10.1.

##### C. 3D surface segmentation in Imaris

1. Open the reconstructed Airyscan image stack in Imaris Bitplane 10.1.
2. Confirm that the individual fluorescence channels are preserved as separate channels.
3. Before creating receptor surfaces, define the analysis volume according to the workflow being performed.
4. For dorsal VNC neuropil analysis, define the analysis volume as the dorsal-most one-third of the acquired VNC z-stack. Begin at the dorsal edge of the image volume and include consecutive optical sections through one -third of the total stack depth. In the image stacks used here, this typically corresponded to approximately 40–45 optical sections. This standardized dorsal volume samples the neuropil where premotor neuron (PMN)–MN and PMN–PMN synapses are concentrated.
5. For MN dendrite-restricted analysis, use a standardized 3D ROI measuring 82.5 µm × 52.5 µm × 7.2 µm. Position the ROI so that it captures the DenMark-labeled MN dendritic territory selected for analysis. Use the same ROI dimensions and placement rule for all samples included in the same comparison.
6. In the *Surpass* workspace, click the blue *Add New Surface* button to begin creating a new surface object. ***Note:*** *For elongated labeled objects, use 3D Surfaces instead of Spots. Surfaces preserve the objects’’ shape and spatial extent, allowing their spatial overlap with other segmented objects to be measured*.
7. In the surface-creation wizard, select *Classify Surfaces* and *Object-Object Statistics. Classify Surfaces* permits later separation of overlapping and non-overlapping receptor objects. *Object-Object Statistics* makes the *Overlapped Volume to Surfaces* measurement available for comparisons between receptor or dendrite surfaces.
8. Click the forward blue arrow to proceed to the segmentation setup page.
9. Under *Source Channel*, select the fluorescence channel containing the first receptor subunit.
10. Under *Method*, select *Background Subtraction (Local Contrast)*.
11. Enter the diameter of the largest sphere that fits within the object. ***Note:*** *To determine the diameter of the largest sphere that fits into a receptor object, open the image in slice view and identify representative receptor puncta in the source channel. Manually measure the diameter of an object in the image plane and use this measurement to select an appropriate sphere diameter for local -contrast background subtraction. Apply the same sphere-diameter setting to all images included in the same quantitative comparison*.
12. Click the forward blue arrow to proceed to the thresholding page.
13. Inspect the intensity histogram and the surface preview alongside the fluorescence image.
14. Adjust the threshold so that the preview follows visible receptor signal while excluding diffuse background.
15. Check several areas of the image, including bright and dim puncta, before proceeding.
16. Under *Split touching Objects (Region Growing)*, select *Enable*.
17. Closely packed receptor puncta can touch in the thresholded image and appear as one continuous surface. Enable this option to separate such connected signal into individual objects for counting and overlap analysis.
18. To select an appropriate Seed Points Diameter, switch to Slice view and examine several clearly isolated receptor puncta in the source channel.
19. Measure the approximate width of individual puncta across their shorter axis, rather than their full length.
20. Enter a diameter representative of these measurements, then inspect the seed-point and surface previews in both isolated and crowded regions.
21. Record the diameter and apply it consistently to images in the same quantitative comparison. ***Note***: A diameter that produces several seeds within one visibly continuous punctum may oversplit it; a diameter that fails to place separate seeds in distinguishable touching puncta may leave them merged.
22. Select *intensity-based splitting*. This setting uses differences in fluorescence intensity within connected signal to guide separation around the seed points.
23. Compare the resulting surface boundaries with the fluorescence image before proceeding.
24. Click the forward blue arrow to proceed to the *Filter Seed Points* page.
25. On the *Filter Seed Points* page, click *Add* and select *Intensity Mean* as the filter type.
26. Confirm that the measurement refers to the intended receptor channel.
27. Inspect the seed-point intensity histogram together with the fluorescence image and surface preview.
28. Adjust the lower threshold to remove seeds arising from background signal while retaining seeds located within visible receptor puncta.
29. Check several areas of the image, including regions with dim puncta, before accepting the threshold.
30. Click the forward blue arrow to proceed to the *Filter Surfaces* page.
31. Click *Add* and select *Number of Voxels* as the filter type. This filter removes small, segmented objects that are likely to represent residual background or noise rather than receptor puncta.
32. Choose the lower voxel cutoff by examining the surface preview alongside the fluorescence image.
33. Identify the smallest clearly visible receptor puncta that should remain in the analysis, then set the cutoff low enough to retain those objects while excluding smaller objects without corresponding receptor signal.
34. Inspect the result across several image regions before accepting the cutoff.
35. Record the selected voxel cutoff and apply the same criterion to images included in the same quantitative comparison, provided they were acquired and reconstructed at the same voxel dimensions.
36. If voxel dimensions differ, reassess the cutoff because a given number of voxels represents a different physical volume.
37. Click the green forward arrow to complete surface creation.
38. nspect the completed surfaces against the fluorescence image to confirm that visible puncta are represented and that obvious background objects have been excluded.
39. Rename the completed surface object to identify the receptor subunit and fluorescence channel.
40. Repeat the surface-creation workflow for the second receptor subunit, selecting its own fluorescence channel as the source. Assess its intensity and seed-point thresholds against that channel’‘s image rather than copying the first channel’‘s intensity values.
41. For MN dendrite-restricted analysis, create a separate 3D surface from the DenMark channel within the standardized MN region of interest. Inspect this surface against the DenMark image to confirm that it represents the dendritic territory used for analysis. Keep the dendrite and receptor surfaces as separate objects for the subsequent overlap filters.

#### D. Calibration of the colocalization overlap criterion using biological controls

1. Generate surfaces for the positive control, α6::HA and α6::OLLAS, and the negative control, Rdl::HA and α5::OLLAS, using the surface-creation workflow in Section C.
2. For each control pair, measure *Overlapped Volume to Surfaces* in both directions and inspect the resulting overlap values alongside the images. Select a cutoff that retains overlap between the two labels of the positive control while excluding the largely separate objects in the negative control.
3. Use ≥0.05 µm^3^ as the criterion for classifying a surface as overlapping.
4. Obtain the number of puncta for each label from the *Statistics* tab in Imaris, using the total number of surfaces for the corresponding channel.
5. This control-defined cutoff produced greater than 98% colocalization for the positive control and less than 2% for the negative control in the published analysis.
6. Apply the same ≥0.05 µm^3^ criterion throughout the experimental analyses.

***Note***: *Establish the overlap criterion before classifying experimental receptor pairs. Keep the selected criterion consistent across images included in the same comparison*.

#### E. Quantification of nAChR subunit colocalization in the dorsal VNC neuropil

1. Select the first receptor surface object generated from the dorsal-most one-third of the VNC z-stack. In the surface filter settings, select *Overlapped Volume to Surfaces* and choose the second receptor surface object as the comparison.
2. Classify puncta with an overlap value of ≥0.05 µm^3^ as colocalized and puncta below 0.05 µm^3^ as non-colocalized.
3. Obtain the number of puncta for each label from the *Statistics* tab in Imaris, using the total number of surfaces for the corresponding channel.
4. Repeat the measurement with the receptor subunits reversed.

#### F. MN dendrite-restricted colocalization analysis

1. Select the surface object for subunit 1. Apply the *Overlapped Volume to Surfaces* filter using the DenMark-labeled MN dendrite surface as the comparison. Retain only subunit 1 puncta with dendrite overlap ≥0.05 µm^3^.
2. Apply the same dendrite-overlap filter to subunit 2 and retain only its puncta with dendrite overlap ≥0.05 µm^3^. Record the number of dendrite-associated puncta retained for each subunit.
3. Apply *Overlapped Volume to Surfaces* again, this time comparing the retained subunit 1 puncta with the retained subunit 2 puncta. Classify subunit 1 puncta with receptor–receptor overlap ≥0.05 µm^3^ as colocalized.
4. Repeat the receptor–receptor comparison in the opposite direction to classify the retained subunit 2 puncta.
5. Obtain the number of puncta for each label from the Statistics tab in Imaris, using the total number of surfaces for the corresponding channel.

***Note***: *The first overlap filter selects puncta associated with the labeled MN dendritic compartment. The second measures overlap between the two receptor subunits within that retained population*.

## Data analysis

1. Export the numbers of colocalized and non-colocalized puncta for each receptor subunit from Imaris. Keep the two directions of each receptor-pair comparison in the data record. For MN dendrite-restricted analysis, also record the number of puncta retained after filtering each subunit against the DenMark-labeled dendrite surface.
2. Calculate the colocalization percentage for each subunit in each image: Colocalization (%) = 100 × [colocalized puncta ÷ (colocalized puncta + non-colocalized puncta)]. For dorsal neuropil analysis, the denominator is all retained puncta of that subunit in the selected dorsal volume. For MN dendrite-restricted analysis, the denominator includes only puncta of that subunit retained by the dendrite-overlap filter. Do not combine results from the two anatomical compartments.
3. Use one VNC from one larva as a biological replicate. Multiple puncta within a VNC are objects used to calculate that replicate’‘s percentage; they are not independent biological replicates. If more than one image or region is analyzed from the same VNC, treat those measurements as technical replicates and summarize them to one value per VNC before comparing groups.
4. Report the colocalization percentage for each VNC together with its underlying puncta counts. Calculate percentages separately for both subunits in a pair. In the published study, these directional percentages were nearly identical, so one representative subunit per pair was used for graphical display and statistical analysis while both measurements were retained.
5. Use ZEN Blue for 3D Airyscan processing and Imaris for surface segmentation, filtering, and object-based overlap measurements. Organize the exported counts and calculate image-level percentages in a spreadsheet or statistical program such as Excel, R, MATLAB, or Python. The analysis requires experience inspecting multichannel 3D images and checking that segmented surfaces correspond to visible structures; expertise in Linux or programming is not required.

## Validation of protocol

This protocol (or parts of it) has been used and validated in the following research article(s): Sitaula et al. (2026). Motor neurons integrate cholinergic inputs through spatial organization of diverse nicotinic receptors. PNAS Nexus (Figure 5).

## General notes and troubleshooting

### General notes

1. Use the same acquisition and surface-generation criteria for images included in a quantitative comparison. Keep the objective, zoom, z-step, channel configuration, and relevant detector settings consistent.
2. Inspect segmentation in both Slice and 3D views. A surface should follow the visible receptor signal without extending into diffuse background or omitting clearly labeled structures.
3. Interpret “Overlapped Volume to Surfaces” using the control-defined cutoff described in the Procedure. The ≥0.05 µm^3^ criterion classifies spatial overlap between segmented objects; it does not establish that two subunits are assembled into the same receptor complex. Check the positive and negative controls when establishing the analysis conditions, particularly after changing staining, imaging, or segmentation settings.
4. Keep voxel dimensions consistent across images being compared and record the minimum-voxel cutoff used to remove small surfaces. If voxel dimensions differ between images, the same voxel cutoff will represent different physical object sizes.
5. Count each independently dissected larval VNC as one biological replicate. Multiple z-slices, image regions, or receptor surfaces measured from that VNC are not additional biological replicates.

### Troubleshooting

Problem 1: Few receptor surfaces are detected, or the surfaces omit visible receptor signal.

Possible cause: Weak or uneven immunolabeling, inappropriate channel selection, incomplete Airyscan reconstruction, or a segmentation threshold that excludes genuine signal.

Solution: Confirm the identity of the source channel and inspect the reconstructed stack alongside the original fluorescence image. Check staining quality across the z-stack before changing the Imaris settings. Revisit the background-subtraction sphere diameter and threshold while viewing several bright and dim regions in Slice view. Adjust the threshold only when the resulting surfaces more faithfully follow visible signal; record the final value and apply the same decision rule across comparable images.

Problem 2: Adjacent receptor objects remain joined, or a single elongated object is divided into many surfaces.

Possible cause: The threshold or seed-point diameter is unsuitable for the size and spacing of the labeled objects, or the seed-intensity filter retains too many or too few seeds.

Solution: Examine isolated and crowded objects in Slice view. Measure the shorter-axis width of several clearly separated objects and use those observations to choose a seed-point diameter. Inspect the seed-point preview and the resulting surfaces before completing creation. Retain splitting only when it separates visibly distinct objects without fragmenting continuous signal, and document the settings used for the comparison.

Problem 3: Many small background objects remain after surface creation.

Possible cause: Diffuse background was included during thresholding, the seed-intensity cutoff is too permissive, or the minimum-size filter is too low.

Solution: Compare the candidate surfaces with the underlying fluorescence image. Adjust the threshold, seed-intensity filter, and Number of Voxels cutoff so that small background objects are excluded while credible receptor objects are retained. After surface creation, inspect the remaining objects in Slice and 3D views and delete individual surfaces that clearly represent background or imaging artifacts. Apply the same criteria to all comparable images and record any manual deletions. Do not remove a surface because of its effect on the colocalization result.

## Supplementary information

No additional supplementary files accompany this protocol. The image stacks and analysis code associated with the original study are available through the Dryad repository cited in the Software and datasets section (doi: 10.5061/dryad.fxpnvx17f).

## Acknowledgments

This study was supported by NIH-NINDS 1R01NS142268-01A1, the Texas A&M University Division of Research Targeted Proposal Teams (TPT) funding program, Zarin Ascend FY25 and FY26, and Texas A&M College of Arts and Sciences, Zarin STRP FY24 and FY25 grants.This protocol is derived from the original work of Sitaula et al. (2026).

## Author contributions

Ankura Sitaula developed and performed the receptor-colocalization experiments and image analyses and drafted the protocol. Aref Zarin supervised the study and reviewed and edited the protocol.

## Competing interests

The authors declare no competing interests.

## Ethical considerations

This protocol uses *Drosophila melanogaster* larvae and does not involve human participants or vertebrate animals.

---

**Figure 1.**
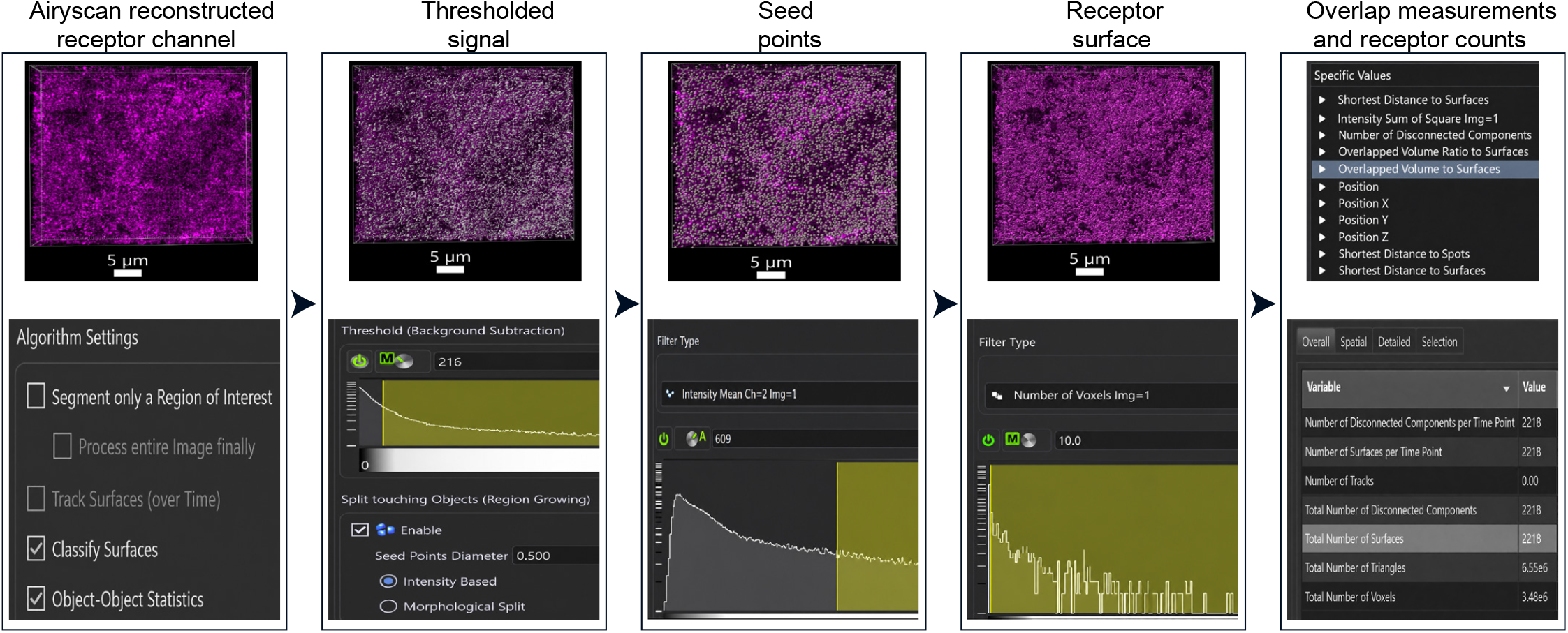
Imaris workflow for 3D receptor-surface segmentation. A reconstructed receptor fluorescence channel is converted into discrete 3D receptor-punctum surfaces for object-based colocalization analysis. From left to right, the panels show the Airyscan-reconstructed receptor channel, thresholded signal, seed points used to separate touching puncta, final receptor surfaces, and selection of the overlap statistic followed by counting of overlapping receptor puncta. The corresponding Imaris settings are shown beneath each image. Thresholding retains receptor-associated signal while excluding diffuse background; seed points are filtered by mean intensity, and resulting surfaces are filtered by voxel count to remove small background objects. The final panel shows selection of the “Overlapped Volume to Surfaces” statistic and the Statistics table, where “Total Number of Surfaces” reports the number of overlapping receptor puncta retained after applying the control-defined cutoff of ≥0.05 µm^3^. The total number of receptor puncta is obtained from the original surface object before applying this filter. The number of non-overlapping puncta is calculated by subtracting the overlapping count from this original total. The segmentation settings displayed in the Imaris screenshots are examples for demonstration only and should not be copied directly for other experiments. Users must determine appropriate segmentation thresholds, seed-point diameters, seed-intensity thresholds, and minimum voxel cutoffs from their own images, verifying that segmentation retains visible receptor puncta while excluding background and avoiding excessive splitting or merging. Apply the selected criteria consistently within each quantitative comparison and reassess voxel -count cutoffs if voxel dimensions change. Scale bars, 5 µm.

**Figure 2.**
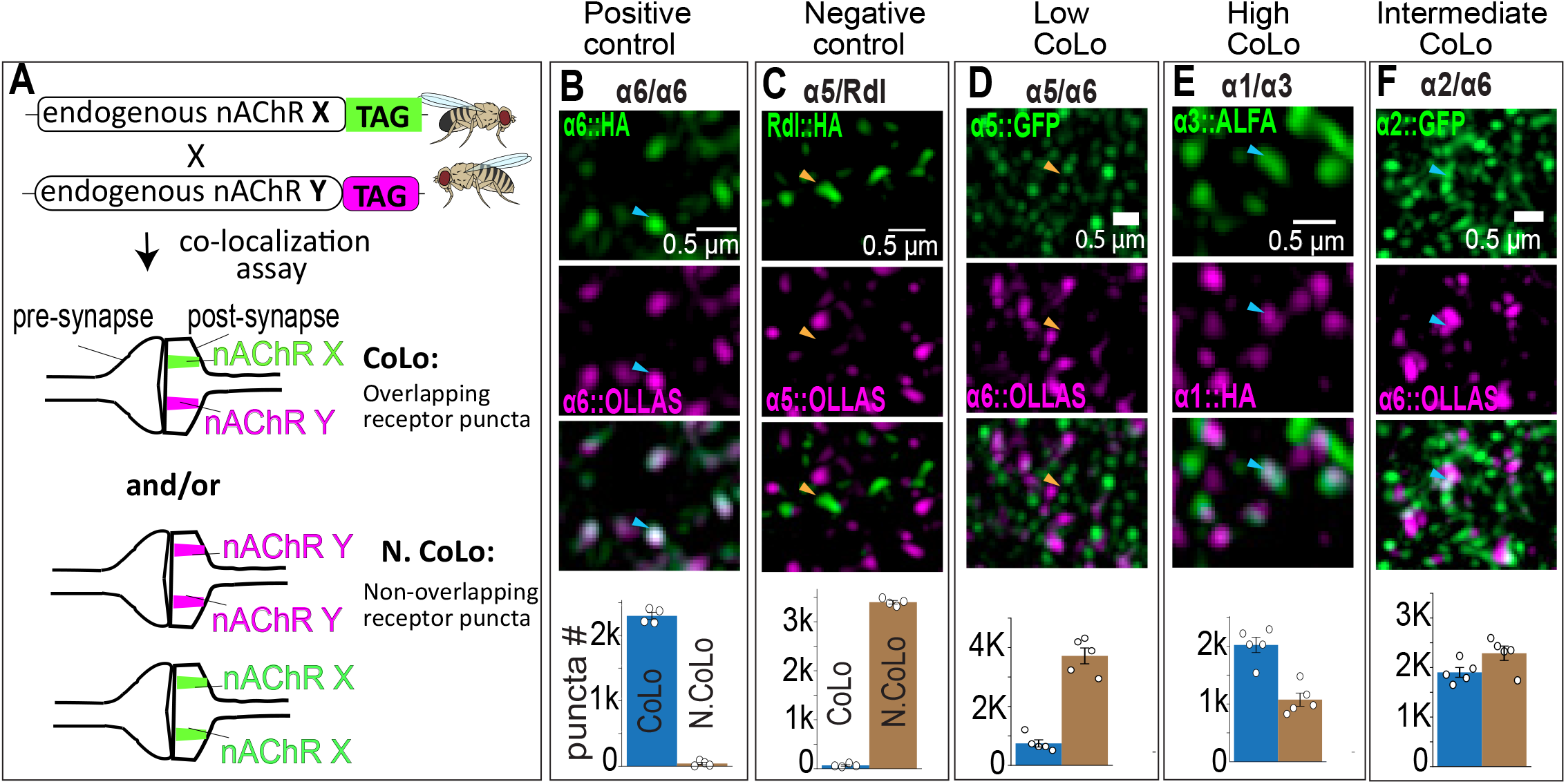
Object-based colocalization and spatial organization of nAChR subunits. (A) Schematic of the assay. Endogenously tagged nAChR subunits X and Y are imaged as separate signals. Puncta are classified as overlapping (CoLo) when the two subunit surfaces meet the control-defined overlap criterion of ≥0.05 µm^3^, or as non-overlapping (N.CoLo) when they do not. (B) Positive control using independently tagged α6::HA (green) and α6::OLLAS (magenta), which show >98% overlap. (C) Negative control using the γ-aminobutyric acid receptor subunit Rdl::HA (green) and α5::OLLAS (magenta), which show <2% overlap. (D) Low-overlap pair α5::GFP (green) and α6::OLLAS (magenta; ∼17% colocalization). (E) High-overlap pair α3::ALFA (green) and α1::HA (magenta; >65% colocalization). (F) Intermediate-overlap pair α2::GFP (green) and α6::OLLAS (magenta; ∼45% colocalization). Panels B–F show each fluorescence channel and the merged image. Blue bars represent overlapping (CoLo) receptor puncta, and brown bars represent non-overlapping (N.CoLo) receptor puncta; this color key applies to panels B–F. The reference populations counted in the bar plots are α6::HA in B, α5::OLLAS in C, α5::GFP in D, α1::HA in E, and α2::GFP in F. CoLo and N.CoLo counts represent puncta of the reference population classified by their overlap with the comparison population. Each dot represents the count in the corresponding category from one independent larval VNC. Bars show the mean, and error bars indicate SEM. Sample sizes are n = 4 independent larval VNCs for panels B and C and n = 5 for panels D–F. Cyan arrowheads indicate representative colocalized receptor puncta, whereas yellow arrowheads indicate representative non-colocalized receptor puncta. Scale bars, 0.5 µm. GFP, green fluorescent protein; HA, hemagglutinin tag.

**Figure 3.**
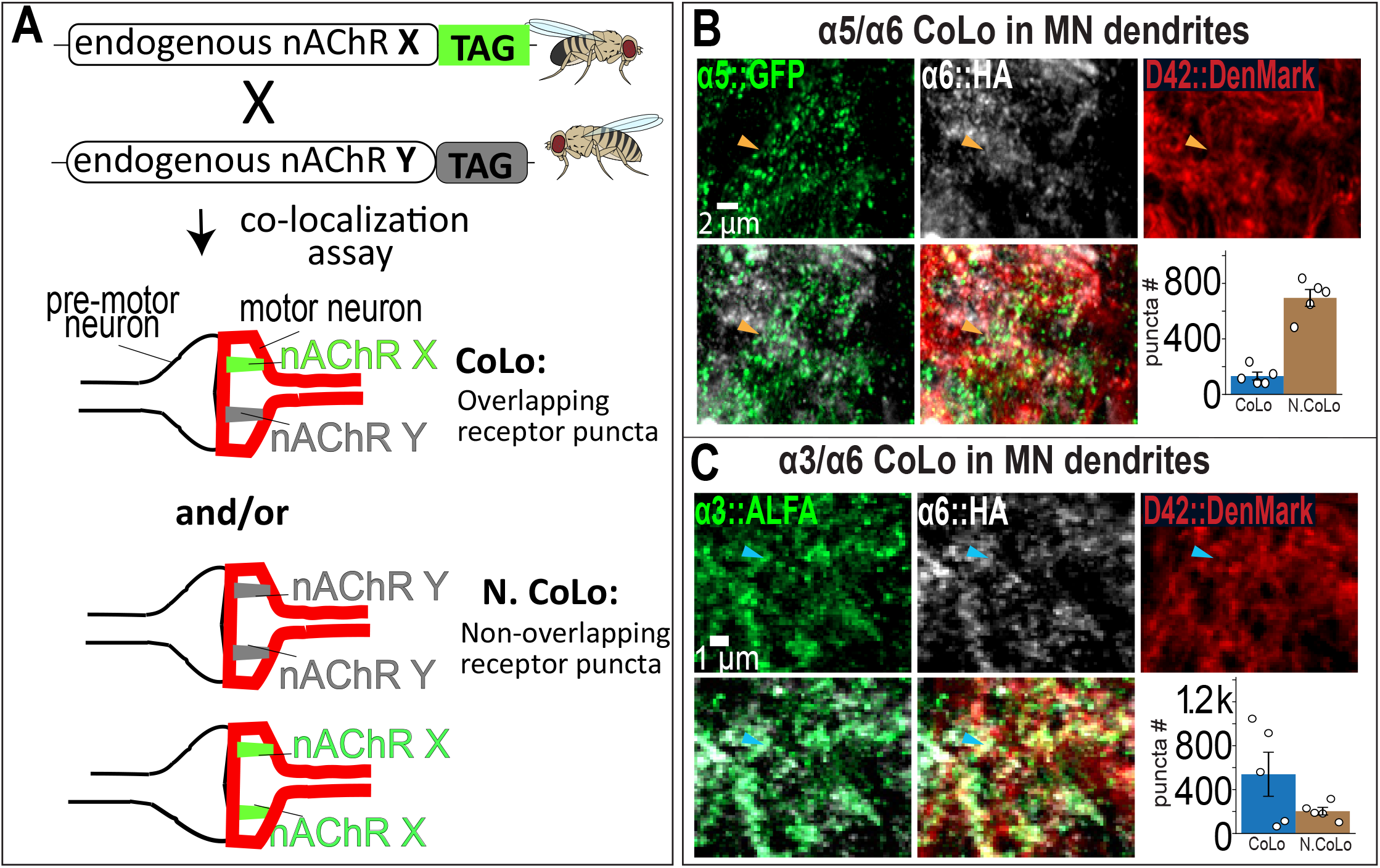
Motor-neuron dendrite-restricted colocalization of nAChR subunits. (A) Schematic of the analysis. Endogenously tagged nAChR subunits X and Y are imaged together with MN dendrites. Receptor puncta are first selected by their overlap with the MN dendritic surface. Within this dendrite-associated population, puncta are classified as overlapping (CoLo) when the two subunit surfaces meet the control-defined overlap criterion of ≥0.05 µm^3^, or as non-overlapping (N.CoLo) when they do not. (B) Low colocalization of α5::GFP (green) and α6::HA (white) within D42::DenMark-labeled MN dendrites (red; approximately 16% colocalization). (C) High colocalization of α3::ALFA (green) and α6::HA (white) within D42::DenMark-labeled MN dendrites (red; approximately 73% colocalization). Panels B and C show the individual fluorescence channels, merged images, and numbers of CoLo and N.CoLo MN-dendrite-associated puncta. Blue bars represent overlapping (CoLo) receptor puncta, and brown bars represent non-overlapping (N.CoLo) receptor puncta. The reference populations counted are α5::GFP in B and α3::ALFA in C, each classified by its overlap with α6::HA. The D42::DenMark mask represents a population of labeled MNs, and the analysis is restricted to their labeled dendrites. Each dot represents the count in the corresponding category from one independent larval VNC. Bars show the mean, and error bars indicate SEM. Sample sizes are n = 5 independent larval VNCs for panels B and C. Cyan arrowheads indicate representative colocalized receptor puncta, whereas yellow arrowheads indicate representative non-colocalized receptor puncta. GFP, green fluorescent protein; HA, hemagglutinin tag. Scale bars, 2 µm in B and 1 µm in C.

## Notes

https://doi.org/10.5061/dryad.fxpnvx17f

### Competing Interest Statement

The authors have declared no competing interest.

## References

1. Kondo, S., Takahashi, T., Yamagata, N., Imanishi, Y., Katow, H., Hiramatsu, S., Lynn, K., Abe, A., Kumaraswamy, A. and Tanimoto, H. (2020). Neurochemical Organization of the Drosophila Brain Visualized by Endogenously Tagged Neurotransmitter Receptors. Cell Reports 30(1): 284-297.e285. 10.1016/j.celrep.2019.12.018.

2. Sanfilippo, P., Kim, A. J., Bhukel, A., Yoo, J., Mirshahidi, P. S., Pandey, V., Bevir, H., Yuen, A., Mirshahidi, P. S., Guo, P., et al. (2024). Mapping of multiple neurotransmitter receptor subtypes and distinct protein complexes to the connectome. Neuron 112(6): 942-958.e913. 10.1016/j.neuron.2023.12.014.

3. Sitaula, A., Kaur, K., Mogharrabi, A., Olsen, L. and Zarin, A. (2026). Motor neurons integrate cholinergic inputs through spatial organization of diverse nicotinic receptors. PNAS Nexus 5(6): pgag173. 10.1093/pnasnexus/pgag173.

4. Rosenthal, J. S. and Yuan, Q. (2021). Constructing and Tuning Excitatory Cholinergic Synapses: The Multifaceted Functions of Nicotinic Acetylcholine Receptors in Drosophila Neural Development and Physiology. Front Cell Neurosci 15: 720560. 10.3389/fncel.2021.720560.

5. Le Novère, N., Corringer, P. J. and Changeux, J. P. (2002). The diversity of subunit composition in nAChRs: evolutionary origins, physiologic and pharmacologic consequences. J Neurobiol 53(4): 447–456. 10.1002/neu.10153.

6. Corringer, P. J., Le Novère, N. and Changeux, J. P. (2000). Nicotinic receptors at the amino acid level. Annu Rev Pharmacol Toxicol 40: 431–458. 10.1146/annurev.pharmtox.40.1.431.

7. Dupuis, J., Louis, T., Gauthier, M. and Raymond, V. (2012). Insights from honeybee (Apis mellifera) and fly (Drosophila melanogaster) nicotinic acetylcholine receptors: from genes to behavioral functions. Neurosci Biobehav Rev 36(6): 15531564. 10.1016/j.neubiorev.2012.04.003.

8. Bolte, S. and Cordelières, F. P. (2006). A guided tour into subcellular colocalization analysis in light microscopy. J Microsc 224(Pt 3): 213–232. 10.1111/j.1365-2818.2006.01706.x.

9. Dunn, K. W., Kamocka, M. M. and McDonald, J. H. (2011). A practical guide to evaluating colocalization in biological microscopy. Am J Physiol Cell Physiol 300(4): C723–742. 10.1152/ajpcell.00462.2010.

10. Costes, S. V., Daelemans, D., Cho, E. H., Dobbin, Z., Pavlakis, G. and Lockett, S. (2004). Automatic and quantitative measurement of protein-protein colocalization in live cells. Biophys J 86(6): 3993–4003. 10.1529/biophysj.103.038422.

11. Rosenthal, J. S., Yin, J., Lei, J., Sathyamurthy, A., Short, J., Long, C., Spillman, E., Sheng, C. and Yuan, Q. (2021). Temporal regulation of nicotinic acetylcholine receptor subunits supports central cholinergic synapse development in Drosophila. Proc Natl Acad Sci U S A 118(23). 10.1073/pnas.2004685118.

12. Samara, E., Schilling, T., Ribeiro, I. M. A., Haag, J., Leonte, M.-B. and Borst, A. (2025). Columnar cholinergic neurotransmission onto T5 cells of Drosophila. Current Biology 35(6): 1269-1284.e1266. 10.1016/j.cub.2025.02.004.

